# Proteasome Substrate Capture and Gate Activation by *Mycobacterium tuberculosis* PafE

**DOI:** 10.1101/249060

**Authors:** Kuan Hu, Jordan B. Jastrab, Susan Zhang, Amanda Kovach, Gongpu Zhao, K. Heran Darwin, Huilin Li

## Abstract

In all domains of life, proteasomes are gated chambered proteases that require opening by activators in order to facilitate protein degradation. Twelve proteasome accessory factor E (PafE) monomers assemble into a single, dodecameric ring to promote proteolysis that is required for the full virulence of the human bacterial pathogen *Mycobacterium tuberculosis*. While the best characterized proteasome activators use ATP to deliver proteins into a proteasome, PafE does not require ATP. In order to understand the mechanism of PafE-mediated protein targeting and proteasome activation, we studied the interactions of PafE with native substrates, including a newly identified proteasome substrate, Rv3213c, and with proteasome core particles. We characterized the function of a highly conserved feature conserved in bacterial proteasome activator proteins: a glycine-glutamine-tyrosine-leucine or "GQYL" motif at their carboxyl-termini that is essential to stimulate proteolysis. Using cryo-electron microscopy, we found that the GQYL motif of PafE interacts with specific residues in the α-subunits of the proteasome core particle to trigger gate opening and degradation. Finally, we found that PafE rings have 40-Å openings lined with hydrophobic residues that form a chamber for capturing substrates prior to the onset of degradation. This result suggests PafE has a previously unrecognized chaperone activity. Collectively, our data provide new insights on the mechanistic understanding of ATP-independent proteasome degradation in bacteria.

---

*Mycobacterium tuberculosis (Mtb*), the causative agent of tuberculosis, has a proteasome system that is essential for causing lethal infections in animals by what are likely to be a multitude of reasons [reviewed in (1)]. In all domains of life, proteasomes include a 20S core particle (CP), which is a barrel-shaped complex formed by stacking of four heptameric rings: two β-rings sandwiched by two α-rings, with some or all of the β-ring subunits having catalytic activity (2-4). In *Mtb*, each β-ring is formed by seven identical PrcB subunits and each α-ring is composed of seven identical PrcA subunits. The *Mtb* 20S CP is tightly gated to prevent unregulated proteolysis (5,6). Proteasomal degradation in *Mtb* is promoted by two distinct pathways [reviewed in (7)]: an ATP-dependent pathway catalyzed by *Mycobacterium* proteasomal ATPase (Mpa) and an ATP-independent pathway mediated by PafE (also known as Bpa for bacterial proteasome activator) (3,8). To stimulate degradation, Mpa or PafE cap either one or both ends of a 20S CP using GQYL motifs at the carboxyl (C)-termini of each monomer of an activator complex. In addition to ATP, the Mpa-proteasome requires proteins to be post-translationally modified with the small protein Pup (prokaryotic ubiquitin-like protein) to target them for degradation (9,10). In contrast, the PafE-proteasome neither requires ATP nor interaction with a post-translational modification on a doomed protein for degradation (8,11,12). Instead, it appears that PafE can facilitate the degradation of unfolded proteins and at least one native protein, HspR, without a post-translational modification (11-13). A failure to degrade HspR by the PafE-proteasome results in slowed growth *in vitro* (12) and likely contributes to the attenuated growth phenotype *in vivo* (11).

PafE is a small protein of 174 amino acids that is functionally similar but evolutionally unrelated to the ATP-independent eukaryotic proteasome activators PA26 and PA28. Like PA26 and PA28, PafE also has a four α-helix bundle fold (13-15). However, PafE is unique in two ways: (1) its unstructured, proteasome-activating C-terminus is 21 amino acids long, which is nearly twice the length of the unstructured C-termini of PA26 and PA28; and (2) PafE monomers assemble into dodecameric rings with 12-fold symmetry (13,16), in contrast to the heptameric rings of PA26 and PA28 (17). We previously investigated the function of the extended PafE C-terminus and found it confers suboptimal binding to 20S CPs; shortening the C-terminus while maintaining the GQYL motif results in a dramatic increase in proteolysis by *Mtb* 20S CPs (13).

The interaction between PafE and the *Mtb* 20S CP was recently studied by cryo-electron microscopy (EM) (16). The structure of the PafE-20S CP complex at a resolution of 3.5 Å shows the asymmetric binding of PafE rings to 20S CPs as well as interactions between the GQYL motifs of PafE and the proteasome α-subunits. However, there are several caveats to the interpretation of this structure. The study used a mutated 20S CP to which PafE would more strongly bind: the amino (N)-terminal eight residues of the α-subunits (PrcA) of the 20S CP were deleted, creating an "open-gate" mutant 20S CP (6); and, the catalytic threonine (Thr) of each 20S CP b-subunit (PrcB) was changed to alanine ("T1A" mutant), which increases binding with PafE (and Mpa and substrates) (11). Thus, the 20S CP was used in a "pre-activated" form, preventing any conclusions from being drawn regarding the impact of PafE binding on proteasome activation. Furthermore, the site-specific cross-linker *para*-benzoyl-L-phenylalanine was used in place of threonine-170 in PafE, a residue immediately preceding the GQYL motif. Conceivably, this cross-linking of PafE with 20S CPs may alternative interactions (16). While these previous studies were informative, we felt that more details on the mechanism of proteasome gate opening by PafE remained to be elucidated.

In this study, we performed cryo-electron microscopy (EM) experiments with wild type (WT) 20S CPs and a gain-of-function PafE mutant to show that PafE requires more than six GQYL motifs to stimulate degradation. Furthermore, the GQYL sequence makes specific interactions with residues in PrcA to cause a rotation of *Mtb* 20S CP α-subunits. This rotation leads to an outward movement and disorder of the PrcA N-termini, which opens the gate into the proteasome core protease. Finally, we found that the large hydrophobic opening of PafE may give it a previously unrecognized chaperone-like activity with the ability to capture substrates for delivery to 20S CPs for destruction.

## RESULTS

### Comparison of the 20S α-ring structures with and without PafE capping reveals a proteasome gate opening mechanism by PafE

We used cryo-EM to understand how PafE with an intact GQYL activation motif opens the substrate entry gate of the *Mtb* 20S CP. PafE rings have a low affinity for 20S CPs, but shortening the C-termini of PafE without deleting the GQYL motifs (PafE_Δ155-166_) increases its affinity for 20S CPs by reducing the distance between a PafE ring and a 20S CP α-ring (13). To better understand the interaction between PafE rings and 20S CPs, we used PafE_Δ155-166_ with 20S CPs for single particle cryo-EM analysis. Roughly 180,000 particles were automatically picked from 4,883 cryo-EM images and then submitted to reference-free 2D classification in RELION2 (18). Only 20S CPs capped by PafE_Δ155-166_ rings on either or both sides were selected for 3D classification (**Supplemental Fig. S1**). Refinement of singly capped 20S CPs without applying any symmetry yielded a 3D density map with an average resolution of 4.2 Å, as estimated by Fourier shell correlation (FSC) of two independent maps calculated with two halves of the particles (**Supplemental Fig. S2** and **Table 1**). However, the local resolution varied considerably: the 20S CP region was in the resolution range of 4 – 6 Å, with resolved secondary structures and even a few large side chains, but the PafE ring had a resolution of ~10 Å. The low resolution of PafE density was expected due to the flexible PafE C-termini interacting with the 20S CPs (13,16). For the doubly PafE-capped 20S CP, we masked PafE densities at both ends of the CP particle, and refined the 20S CP region alone by applying D7 symmetry, which led to an improved 3D density map at 3.4 Å average resolution (**Supplemental Figs. S1, S3A-B**, and **Table 1**). Based on a local resolution estimation, the β-rings of the 20S CP have a better resolution of 2.5 – 3 Å, and the α-rings at the two ends have a lower resolution of 3.2 – 3.8 Å (**Supplemental Fig. S3C**).

**Table 1.** EM data collection and refinement statistics.

| PafE-bound 20S CP |  |  |
| --- | --- | --- |
| Data Collection |  |  |
| Pixel size (Å) | 1.09 |  |
| Under focus (μm) | 1.0 - 3.0 |  |
| High tension (kV) | 300 |  |
| Electron dose (e<sup>-</sup>/Å<sup>2</sup>) | 49 |  |
| Micrographs | 4883 |  |
| 3D reconstruction and refinement |  |  |
|  | Singly PafE-capped 20S CP | Doubly PafE-capped 20S CP |
| Particles | 60,034 | 51,091 |
| Resolution (Å) | 4.2 | 3.4 |
| Symmetry applied | C1 | D7 |
| Model composition |  |  |
| Non-hydrogen atoms | 46500 | 46,618 |
| Protein residues | 6,170 | 6,174 |
| RMSD |  |  |
| Bonds (Å) | 0.007 | 0.007 |
| Angles (°) | 1.21 | 1.21 |
| Dihedrals (°) | 11.83 | 11.83 |
| Ramachandran plot |  |  |
| Favored (%) | 96.9 | 97.21 |
| Allowed (%) | 3.03 | 0.90 |
| Outliers (%) | 0.07 | 0.00 |
| Rotamer outliers | 1.01 | 0.90 |
| MolProbity validation |  |  |
| Clash score | 13.43 | 14.48 |
| Average B-factor | 80.94 | 81.31 |
| Data deposition |  |  |
| EMDB ID | EMD-7098 | EMD-7097 |
| PDB ID | 6BGO | 6GBL |

Although the singly-capped 20S CP structure had a lower resolution than the doubly-capped 20S CP structure, it provided a unique opportunity to compare the open versus closed gates within the same structure (**Fig. 1A**). As expected, the EM map of PafE_Δ155-166_-bound 20S CPs shares a similar architecture with the 3D EM maps reported previously, which were either at a much lower resolution of 12 Å (13), or at a resolution of 3.5 Å but using chemically modified PafE and pre-activated 20S CPs (16). At the PafE-capped end, the first ordered residue in PrcA is Ser-8. A cloud of discontinuous densities in the center and above the α-ring is visible when the 3D map is rendered at a lower display threshold of 5.8s (**Fig. 2B**, left panel). These weak and discontinuous densities are presumably from the disordered amino N-termini of PrcA, each consisting of seven amino acids. The discontinuous densities disappeared when the 3D map was surface rendered at the normal threshold of 7.0σ, revealing an opening of 27 Å in diameter (**Fig. 1B**, right side). The uncapped end of the 20S CP, by contrast, had the gate tightly sealed by well-defined PrcA N-termini (**Fig. 1C**). When the PafE-capped half of the 20S CP was aligned to the uncapped half, we found that the β-rings were unchanged, but each of the PrcA subunits had undergone a 3°, in-plane rotation (**Fig. 1D, Supplemental Fig. S4**). PrcA rotation moves its N-terminal α-helix 0 (H0) outward by 3 Å, thereby increasing the overall diameter of the opening by 6 Å (**Fig. 1D**).

**Figure 1.**
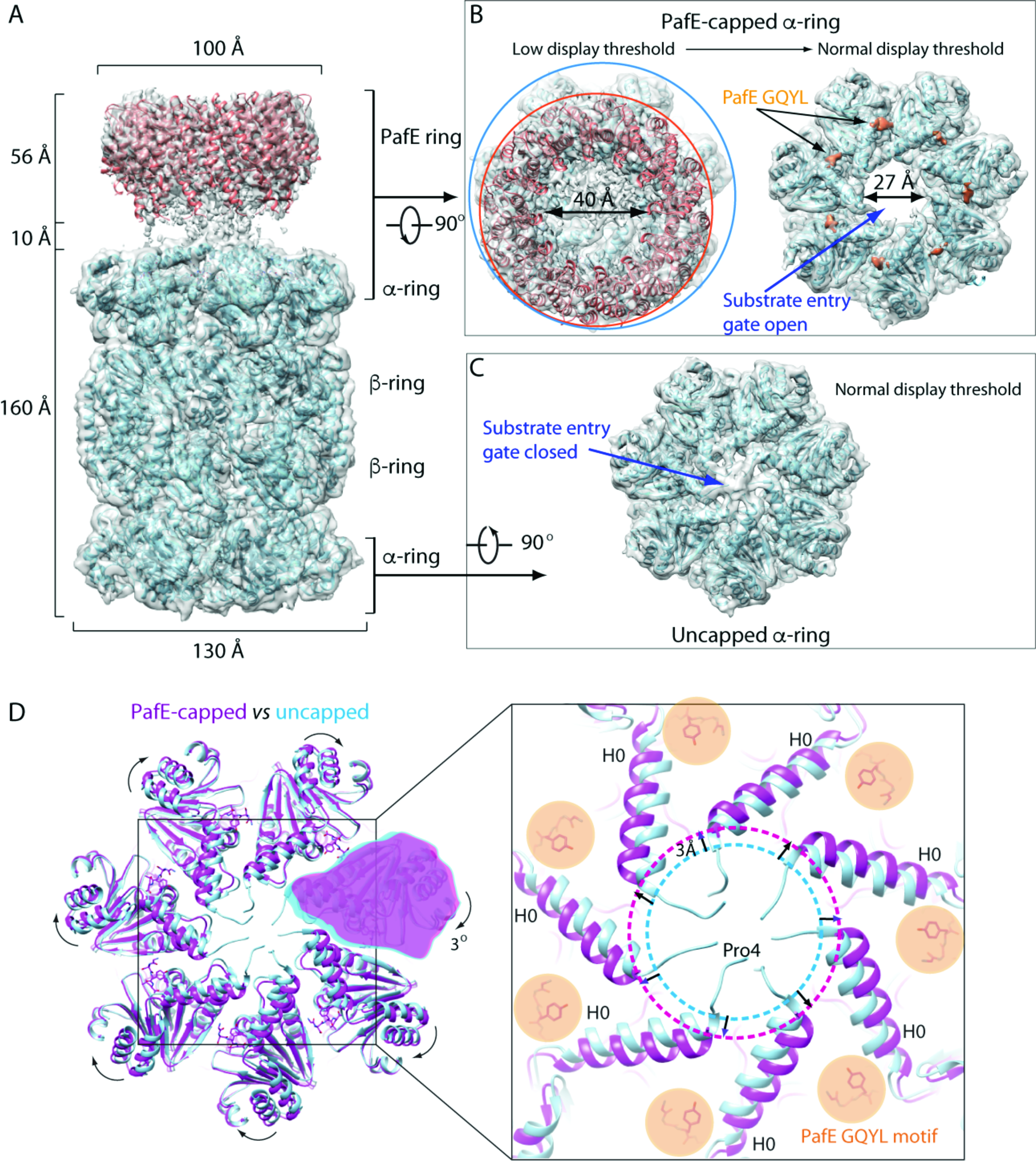
Cryo-EM structure of the PafE_Δ155-166_:20S CP complex. (**A**) Side view of surface-rendered 3D density map of a PafE_Δ155-166_:20S CP complex, shown as a semi-transparent surface, superimposed with the crystal structure of PafE (Salmon, PDB: 5IET) and a refined *Mtb* 20S CP atomic model (cyan, PDB: 3MI0) in cartoons. (**B**) Top view of the PafE_Δ155-166_:20S CP density map surface rendered at a lower threshold (5.8σ) showing the weaker density of PafE ring (left) or rendered at the normal threshold (7.0σ) such that most PafE ring is invisible except for the well-ordered C-termini (right). The densities of PafE C-termini are colored in salmon. The orange and cyan circles mark the circumferences of the PafE and 20S CP, respectively. (**C**) A view of the uncapped end of a 20S CP by 3D density map rendered at the normal threshold (7.0σ). (**D**) Left: Superimposition of the α-rings of the 20S CP with (in magenta cartoon) and without (in cyan cartoon) the capping PafE_Δ155-166_ ring (not shown), revealing a 3° in-plane rotation of each α-subunits. The PafE GQYL motif is shown as orange sticks. Right: a zoomed view of the area marked by the black box in the left panel. Note that the first modeled residue of the α-subunit is Pro-4 and Ser-8 in the uncapped end and PafE-capped end, respectively.

**Figure 2.**
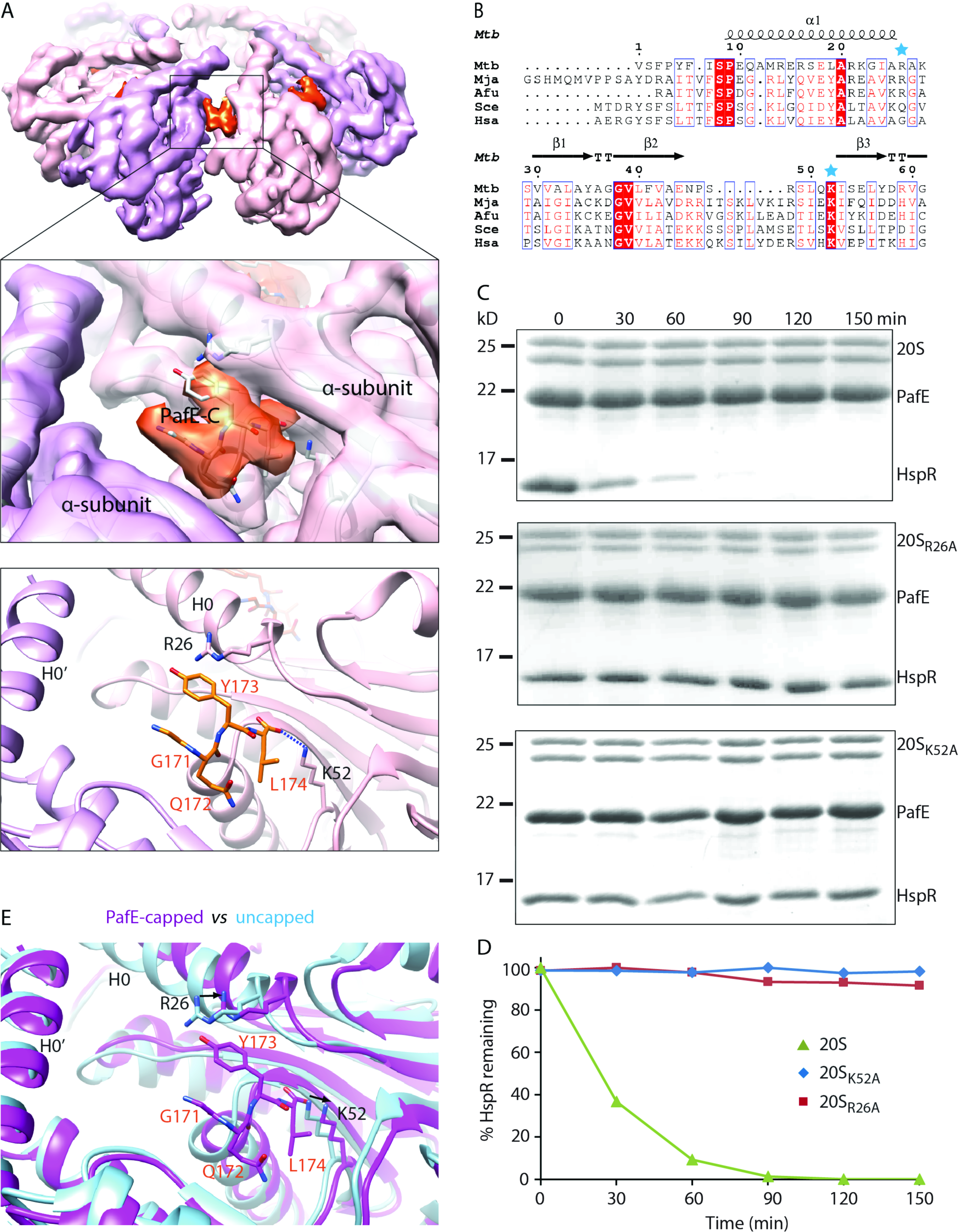
Insertion of the PafE GQYL motifs into the 20S CP moves Helix 0 (H0) of the α-subunit ring. (**A**) Electron density map of the α-ring of the 3.4 Å 3D map of PafE_Δ155-166_-bound 20S CP with D7 symmetry. The PafE C-terminus density is colored in salmon. Middle and lower panels are zoomed-in density and cartoon view of interactions between a PafE C-terminus (salmon) and 20S CP α-subunits (magenta). The side chains of PafE-Y173 and the carbonyl oxygen of PafE-L174 interact with side chains of PrcA-R26 and PrcA-K52, respectively. (**B**) Sequence alignment of the proteasomal α-subunits from different organisms: Mtb; Mja, *Methanococcus jannaschii;* Afu, *Archaeoglobus fulgidus;* Sce, *Saccharomyces cerevisiae;* and Hsa, *Homo sapiens*. In the cases of eukaryotes (Sce and Hsa), the sequences of a2-subunits were chosen for alignment, because these subunits interact with the activating C-termini of the hexameric ATPases Rpt1-6 in the cryo-EM structures of the yeast and human 26S proteasomes (PBD ID 5WVK and 5GJQ). (**C**) SDS-PAGE of in vitro degradation assays using HspR as a substrate. Aliquots were removed at the indicated time points (min) and analyzed by 15% SDS-PAGE. The degradation assays were repeated at least three times with essentially same results. PafE mediates HspR degradation by WT 20S CPs, but not 20S_R26A_ or 20S_K52A_ CPs. (**D**) An quantification of the proteolysis reactions in (**C**) by estimating the relative intensity of each band, showing the amounts of remaining HspR substrate over 150 min reaction time. (**E**) A zoomed in view of superimposed 20S CP α-rings with (magenta) or without (cyan) a capping PafE ring. PafE residues are labeled in orange, and PrcA residues in black. The black arrows show the movements of the side chains in the two structures.

### PafE GQYL motif interacts with R26 and K52 of the 20S CP α-subunits

The 3.4 Å resolution of the doubly PafE-capped 20S CP structure provided a detailed description of the interactions between the activation GQYL motifs of PafE rings with PrcA subunits in the α-ring (**Fig. 2A**). We found Y173 of PafE forms a cation–π interaction with R26 of PrcA, while the carboxyl group of L174 of PafE forms a salt bridge with K52 of PrcA. Amino acid sequence alignment of the 20S α-subunits shows that K52 is conserved across the domains of life, but R26 is conserved only in prokaryotes (**Fig. 2B**). We previously demonstrated K52 of PrcA is essential for PafE-dependent degradation (**13**). To test the significance of the interaction between PafE Y173 and PrcA R26, we substituted R26 to an alanine (PrcAR26A) and tested if the mutated 20S CP could degrade HspR. We found that 20S CPs with PrcAR26A failed to degrade HspR (**Fig. 2C, D**). This result suggests that PrcA R26 is essential for PafE to stimulate gate opening to substrates. Alignment of the atomic models showed both PrcA R26 and K52 were pushed outwards in the presence of the PafE GQYL motif (**Fig. 2E**). Taking these data together, we suggest the mechanism of 20S CP gate opening in *Mtb* is as follows: upon insertion of the PafE C-terminal GQYL motifs into the binding pockets between 20S CP α-subunits, PrcA R26 and K52 move outwards to form a cation–π bond and a salt bridge with the GQYL sequence, which triggers a 3° rotation of the α-subunits, leading to an outward movement of the PrcA H0 helices. This movement disorders the N-terminal gate-lining peptides preceding H0 helices as they lose contact with each other, widening the gate to allow substrates entry into the proteasome core.

### A PafE dodecamer with six GQYL motifs is unable to activate *Mtb* 20S CPs

The consequences of the 12-fold symmetry of a PafE ring are twofold: it leads to a large, 40 Å-wide central channel, nearly twice the diameter of the PA26 and PA28 channels (14,15,17); and it has 12 C-termini that can potentially interact with 20S CPs, nearly twice the number of C-termini provided by PA26 and PA28. Additionally, most proteasome activators have hexameric symmetry, thus we first sought to ask if PafE could use only six of its 12 C-terminal GQYL motifs to stimulate degradation. In order to test this hypothesis, we made a linear PafE dimer ("PafE-PafE"). We cloned DNA encoding a C-terminal-truncated PafE (PafE_1-155_) upstream of a sequence encoding full length PafE (PafE_1-174_), with a linker sequence that encodes a flexible five-amino-acid peptide (GSGNS) in between the two genes (**Fig. 3A**). Complexes of the purified fusion protein eluted from a Superose-6 gel filtration column at the same volume as WT PafE rings (**Supplemental Fig. S5**). Negative stain EM showed that PafE-PafE assembled into rings with a diameter of 10 nm, identical to the WT PafE ring (**Fig. 3A**). These results indicated that we achieved our goal of acquiring a dodecameric PafE ring with only six GQYL-containing termini. We next examined if this variant PafE could activate *Mtb* 20S proteolysis by using HspR (heat shock protein repressor), a known PafE-proteasome substrate in *Mtb* (11). In contrast to the robust degradation of HspR using WT PafE and 20S CPs, PafE-PafE could not degrade HspR (**Fig. 3B**). This result suggests that PafE needs more than six GQYL motifs to effectively activate the degradation of a native substrate by the *Mtb* proteasome.

**Figure 3.**
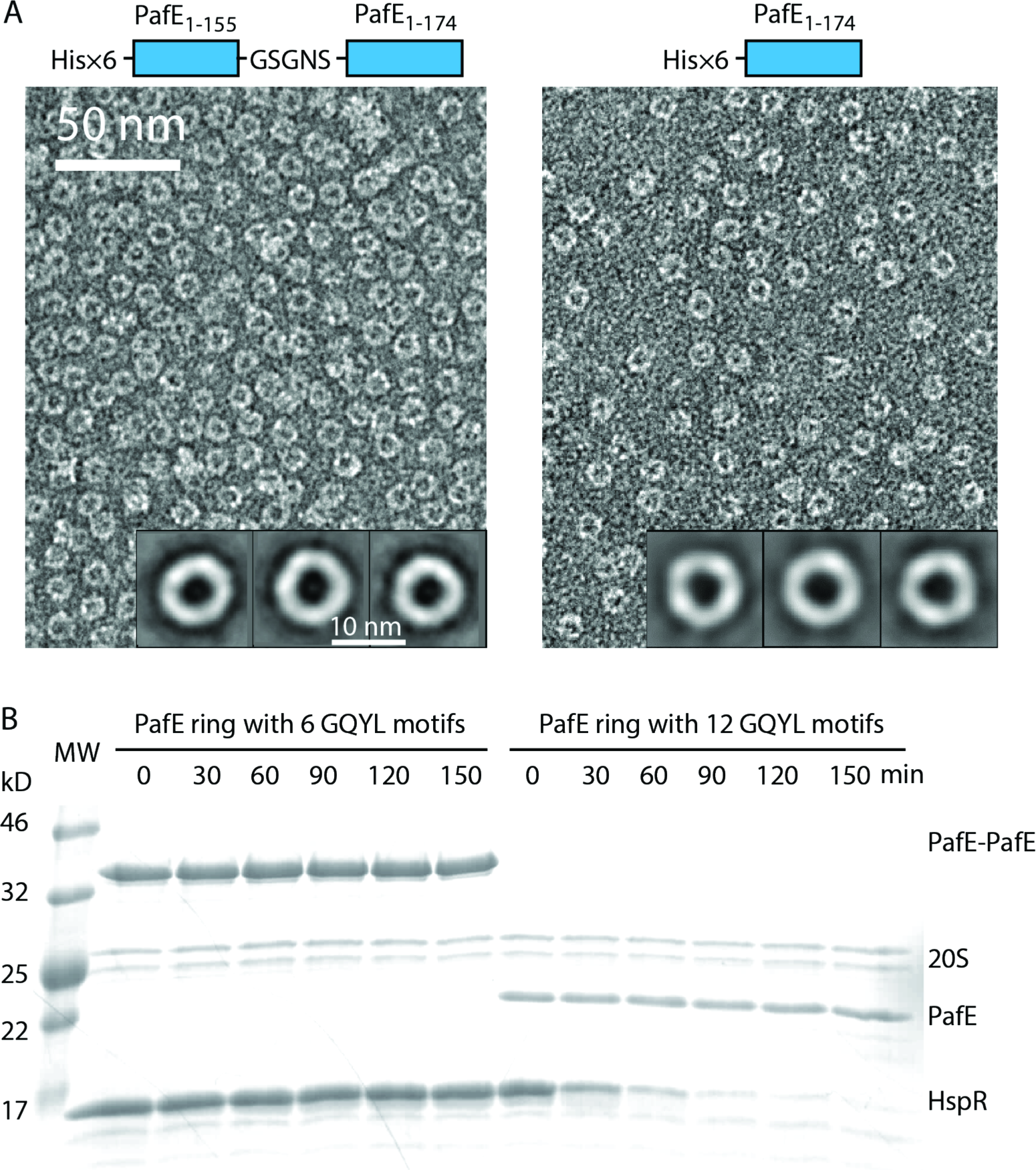
PafE rings with six GQYL motifs cannot mediate substrate degradation by *Mtb* 20S CPs. (**A**) Left: Diagram of the PafE-PafE protein and an electron micrograph of purified and negative stained PafE-PafE protein. Inset: 2D class averages of PafE-PafE. Right: Diagram of wild type PafE and the electron micrograph and 2D class averages of purified protein at the same scale with that of PafE-PafE. (**B**) SDS-PAGE analysis of proteolysis products at the specified time points. Molecular weight (MW) markers are on the left. The degradation assay was done at least three times with essentially same results.

### ParP (Rv3213c) is a PafE-proteasome substrate

Although HspR is the only PafE-mediated proteasome substrate established so far, we previously identified several other proteins that accumulate in an *Mtb pafE* null mutant or in *Mtb* treated with a proteasome inhibitor (11). Furthermore, we also identified numerous proteins that specifically co-purify with a proteasome trap (11). Rv3213c is the only protein that was identified using all three approaches. Rv3213c is predicted to be a ParA-like protein with ATPase activity. ParA, which is conserved in almost all (if not all) bacterial species, works with a partner protein, ParB, and is involved in chromosome partitioning (19). Numerous ParA homologues are present in many bacterial genomes and do not necessarily work with a ParB-like partner protein. *Mtb* has a *bona fide* ParAB pair therefore Rv3213c is not likely to be involved in chromosome partitioning. Furthermore, Rv3213c does not have a conspicuous ParB partner, and it is also not predicted to be essential for viability (20). Because Rv3213c associates strongly with purified proteasomes from *Mtb*, we named it "ParP" for "<u>par</u>titioning with the proteasome".

To test whether or not ParP is a substrate of PafE-mediated degradation, we tried to produce ParP-His_6_ in *E. coli* for *in vitro* degradation assays. However, ParP-His_6_ was insoluble under numerous conditions tested (not shown). We tested if co-production with PafE could help solubilize ParP; remarkably, co-expression of *parP-his*_6_ and *pafE* in *E. coli* resulted in the robust production of soluble ParP-His_6_ (**Fig. 4A**). Importantly, the addition of 20S CPs to the copurified proteins resulted in the robust degradation of ParP-His_6_ (**Fig. 4B, C**). To demonstrate that the ParP degradation was truly mediated by PafE, we truncated PafE to remove the C-terminal 20 residues including the GQYL motif (PafE_Δ155-174_). While ParP could still co-purify with PafE_Δ155-174_ it could no longer be degraded by 20S CPs (**Fig. 4B, C**). Although we cannot rule out the possibility that ParP can be degraded in a PafE-independent manner, our data strongly suggest that ParP is the second, endogenous PafE-proteasome substrate identified in *Mtb*.

**Figure 4.**
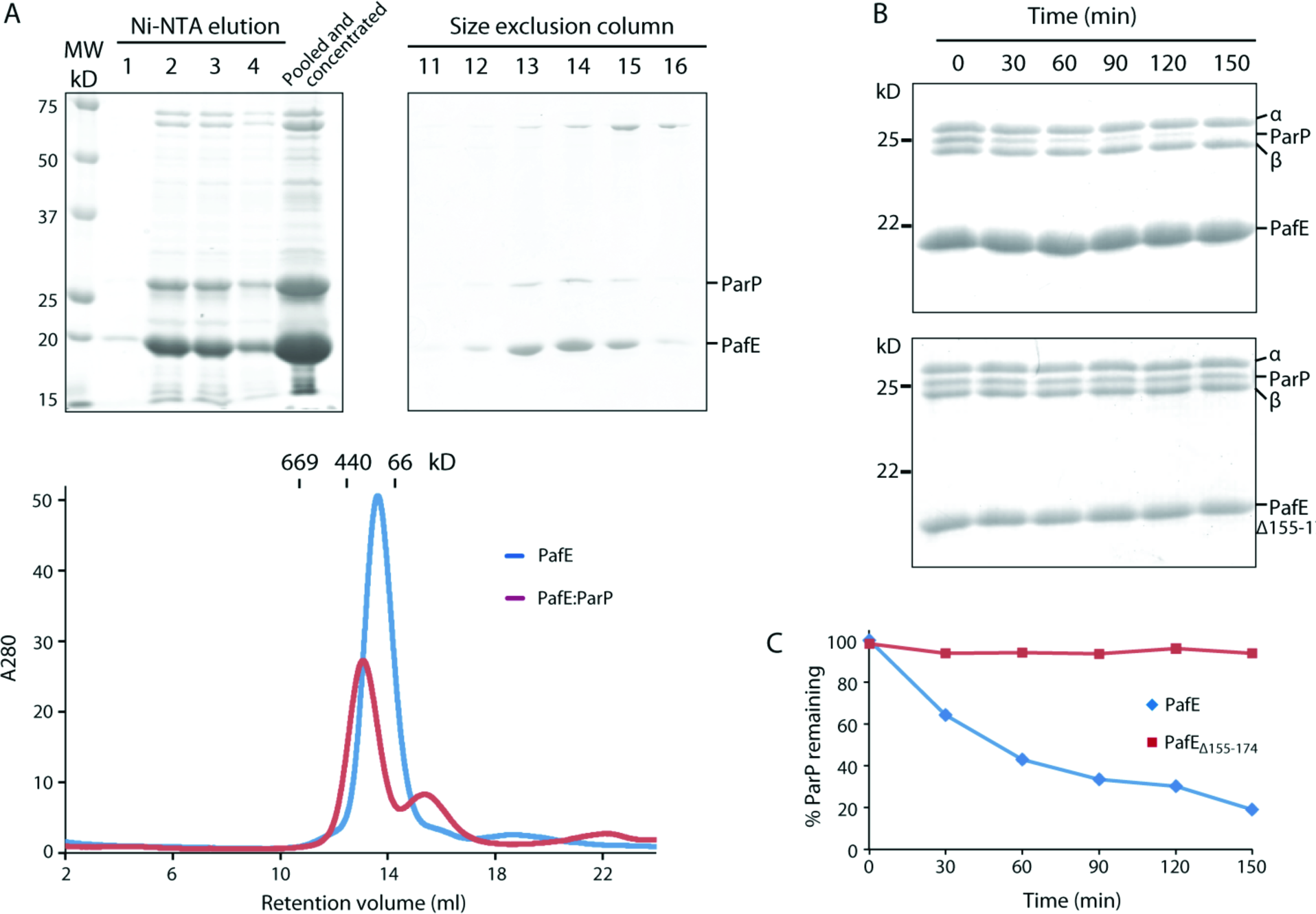
ParP co-purifies with PafE and is degraded by *Mtb* 20S CPs *in vitro*. (**A**) Upper: SDS-PAGE (15% w/v) of ParP-His_6_ co-purified with WT PafE (untagged) as a complex, after Ni-NTA column (left panel) or after further purification with Superose 6 10/300 GL gel filtration column (right panel). Lower: the size exlusion column profile of PafE and PafE:ParP. (**B**) SDS-PAGE (15% w/v) of in vitro proteolysis reaction products. Purified ParP and 20S CPs (both His_6_-tagged) were incubated at room temperature with either PafE (upper panel) or PafE_Δ155-174_ (lower panel). Aliquots were removed at the indicated time points and analyzed. The degradation assays were done at least three times with essentially same results. (**C**) An quantification of the proteolysis reactions in (**B**) by estimating the relative intensity of each band, showing the amounts of remaining ParP substrate over 150 min reaction time.

### PafE substrates are held in a hydrophobic chamber prior to degradation

How an ATP-independent proteasomal activator targets a substrate for degradation by 20S CPs is incompletely understood. Conceivably, the activators could simply bind to the 20S CP and open it for substrate entry, thereby passively allowing the substrate to enter the 20S CP for degradation. However, the finding that PafE has defined substrates such as HspR and ParP suggest otherwise; it is possible that PafE plays a more active role in targeting its substrates for degradation. Given that PafE and ParP form a stable complex, we investigated how ParP associates with the PafE ring by using negative stain EM.

In averaged EM images obtained from reference-free image classification, we found a density or dot in the middle of the PafE ring that was absent in averaged images of PafE without ParP (11) (**Fig. 5A**). Therefore, the dots occupying the PafE channels were most likely ParP. ParP has a molecular weight of 28 kD, potentially small enough to be accommodated inside the 40 Å central channel of a PafE ring.

**Figure 5.**
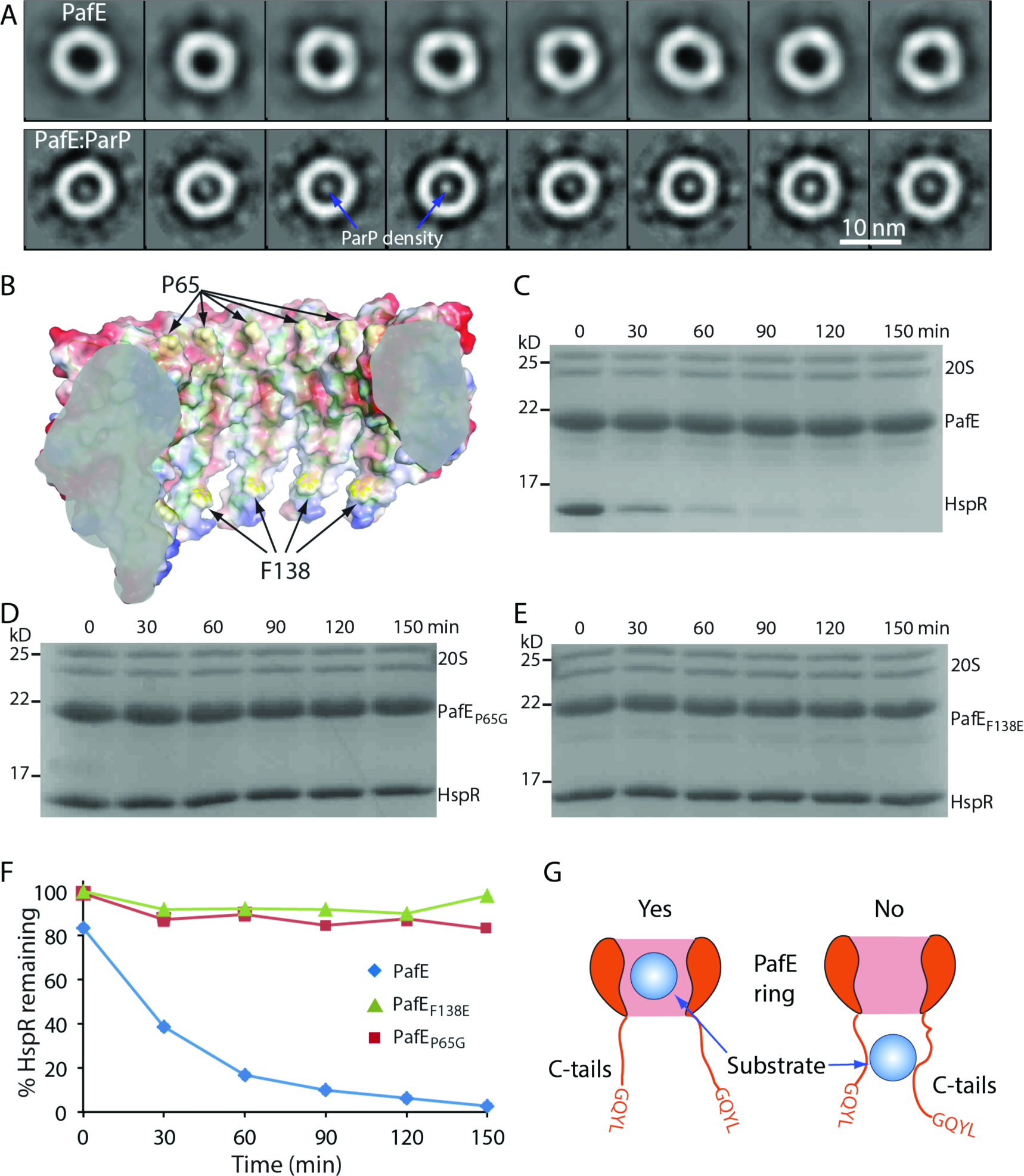
Proteins are targeted for degradation to 20S proteasomes through the central channel of PafE rings. (**A**) Representative reference-free 2D class averages of negatively stained PafE alone (upper panel) and ParP:PafE (lower panel) purified from *E coli*. (**B**) A cut-open side view of PafE ring showing the interior chamber surface. The surface potential map of PafE (from blue positive charge to red negative charge) is superimposed on the cartoon view of the same structure, with the two hydrophobic residues P65 and F138 lining the chamber surface shown in yellow spheres. (**C-E**) SDS-PAGE (15%) of HspR degradation reaction products. Recombinant 20S CP-His_6_ and HspR-His_6_ were incubated at room temperature with either His_6_-PafE (**C**), PafE_P65G_ (**D**), or PafE_F132E_. (**E**) Aliquots were removed at the indicated time points and analyzed. (**F**) Quantification of the remaining HspR over the time course by estimating the relative intensities of individual bands. (**G**) A model whereby substrate is held inside a well-ordered PafE chamber, and not by the C-termini.

In the absence of substrate, the 40 Å central chamber of a PafE ring is likely filled with water. We noticed that the interior surface of the chamber is lined with hydrophobic residues, P65 and F138, which form two hydrophobic belts inside the channel (13). We envisioned that ParP may be held in a PafE ring either by binding to the hydrophobic inner surface of the ring or by binding to a more flexible space formed by the twelve C-terminal, 21 amino acids that form a "beaded curtain". Because we previously found that shortening the C-termini by a few amino acids while keeping the GQYL motif intact does not compromise PafE function and actually enhances protein degradation (13), this finding argues against the possibility that ParP is held by the C-termini.

We mutated the hydrophobic residues P65 and F138 to Gly and Glu, respectively (**Fig. 5B**). Both PafE mutants formed the same oligomers as WT protein based on size exclusion chromatography (**Supplemental Fig. S5**). However, neither of the two mutant PafE alleles formed a complex with ParP (data not shown), indicating that these hydrophobic residues are required for binding to ParP. Because ParP could not be purified in the absence of WT PafE, we used HspR to assess the ability of the mutant PafE alleles to mediate substrate degradation by 20S CPs. As with ParP, the ability of PafE to mediate the proteolysis of HspR was abolished by either single mutation (**Fig. 5C-F**). This result supports the hypothesis that PafE rings must interact with substrates via their structured hydrophobic channel, and not with the flexible 12 C-termini, before delivering proteins into the proteasome core protease (**Fig. 5G**).

## DISCUSSION

Our structural and biochemical studies of the PafE-proteasome system identified specific interactions between PafE and the 20S CP that cause conformational changes leading to the opening of the proteasome gate to substrates. Furthermore, our data suggest a mechanism of PafE-mediated degradation in which a disordered or partially disordered protein is selected by and accommodated within the hydrophobic central channel of a PafE ring. We found that PafE is lined with hydrophobic residues that are essential for mediating proteolysis, and these hydrophobic residues were critical for allowing a robust interaction with a newly identified PafE-proteasome substrate, ParP. Interestingly, this interaction occurred in the absence of proteasomes, suggesting the intriguing possibility that PafE could act as a chaperone to deliver proteins to the proteasome for destruction (**Fig. 6**).

**Figure 6.**
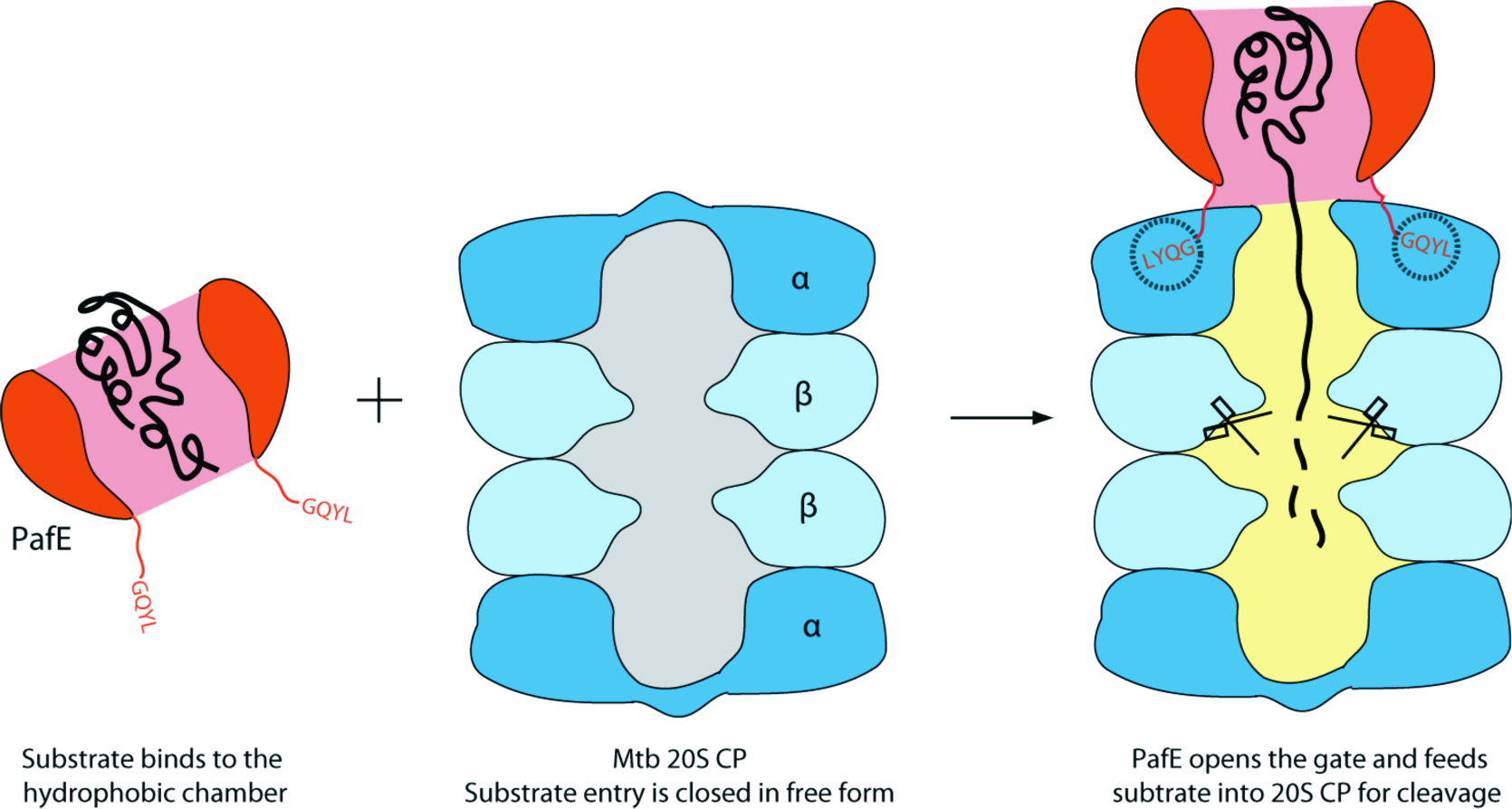
A proposed model for PafE-mediated protein substrate degradation by *Mtb* 20S CP. PafE dodecamer has a central channel with a diameter of 40 Å. The sizable chamber is partially hydrophobic, and accommodates small or partially disordered protein substrates for degradation by the *Mtb* proteasome. The substrate entry gate in the *Mtb* 20S CP is closed in its free form, and is opened by the C-termini of PafE when a PafE ring docks onto the α-ring. The disordered substrate held inside PafE chamber passes through the open gate and reaches the proteolytic site in the β-rings and is subsequently cleaved.

Our cryo-EM structure of the PafE ring with an intact activation motif in complex with a WT 20S CP revealed that the GQYL motif inserts into an acceptor pocket at the interface between two neighboring α-subunits. K52 of the α-ring acceptor pocket is known to play a crucial role by forming a salt bridge with the terminal carboxylate of the PafE activation motif (15,21-23). Importantly, our structure showed an additional residue, PrcA R26, plays a role in gate activation by forming a cation–σ interaction with PafE Y173 of the GQYL motif. These interactions cause an in-plane rigid-body rotation of each of the seven PrcA subunits. The net outcome of these changes is the increase of the gate diameter, leading to a loss of interaction and disorder among the seven gate-forming N-terminal peptides of the α-ring. To our knowledge, this is the first time the gate-opening mechanism of a bacterial 20S CP by an ATP-independent activator has been elucidated.

Structural studies of eukaryotic ATP-independent proteasomal activators showed varied molecular mechanisms of proteasome activation [reviewed in (24)]. *Trypanosoma brucei* PA26 inserts its activating C-terminal between the α-subunits of the *Saccharomyces cerevisiae* 20S CP and induces an outward displacement of a reverse turn of α-subunits to open the gate (14,21). Such a reverse turn is conserved in archaeal and eukaryotic 20S CPs but is absent from the *Mtb* 20S CP (5). In contrast, the monomeric *S. cerevisiae* Blm10 docks a single activation loop between the a5 and a6 subunits of the yeast 20S CP to cause disorder and openning of the gate (25). Consistent with the different activation mechanisms, the last four amino acids of the ATP-independent activators are quite diverse; for example, they are HMVS in *T. brucei* PA26, GMIY in human PA28α, FSMY in human PA28b, and ETLY in human PA28g, as compared to GQYL in PafE. Such variation may have evolved to confer specificity towards their respective 20S CPs. Despite the mechanistic variation, a prominent conserved feature across the domains of life is the formation of salt bridges between the C-terminal carboxylates of the activators and the conserved lysine of the α-subunits of 20S CPs.

Our observation that mutant PafE dodecamers with six GQYL motifs fail to activate 20S CPs suggests that for a PafE ring to engage sufficient 20S CP acceptor sites, it must have more than six GQYL motifs. The 12 activation motifs of a PafE ring are likely required to achieve the full occupancy of the acceptor pockets within an Mtb 20S CP. The term "full occupancy" remains to be defined; an argument against the *Mtb* 20S CP requiring the occupancy of all seven acceptor sites is made by the fact that the ATPase activator Mpa is functional with only six GQYL motifs (26,27). It is further important to note that just because our recombinant PafE-PafE ring has six GQYL motifs, it does not necessarily mean it can engage six acceptor sites in a 20S CP. Taken together, a PafE ring with 12 GQYL motifs may be better able to engage a sufficient number of 20S CP acceptor sites to stimulate degradation.

Despite extensive effort, robust proteasomal activation by Mpa with WT 20S CPs has not yet been demonstrated *in vitro*. This observation may not be surprising in light of our previous studies showing the Mpa GQYL motif is largely hidden inside the substrate translocation channel. Therefore, a large conformational change is likely required for Mpa to bind and activate 20S CPs. These activities may require assistance from additional, yet-to-be-identified co-factor(s) (27). Thus, Mpa and PafE may have additional differences in their mechanisms for activating degradation by 20S CPs. There is also some consensus on ATP-independent and ATP-dependent proteasomal activation in terms of the roles of the 20S α-subunit residues, such as K52 and R26 in *Mtb* PrcA. A lysine is highly conserved at the equivalent position in the α-subunits in all domains of life, and interacts with C-terminal carboxylate groups of ATP-dependent activators, including PAN (28) and Rpts (23,29), as well as ATP-independent activators, including PA26 (14) and Blm10 (25). The cation–σ interaction formed between an activator's penultimate tyrosine and positively charged residue corresponding to *Mtb* PrcA R26 in other proteasome α-subunits has also been observed in the archaeal PAN-20S CP interaction (28); however, this residue (R or K) in both archaeal and bacterial α-subunits, is less well conserved in eukaryotic proteasomes.

An important new finding was the discovery of ParP as a new PafE-proteasome substrate. Remarkably, we found that PafE holds ParP inside its hydrophobic chamber in the absence of 20S CPs. Altering the hydrophobicity of the PafE chamber abolished ParP binding as well as its degradation by 20S CPs. This discovery raises the possibility that the bacterial activator may be able to select and target specific folded substrates to 20S CPs or, alternatively, PafE may bind specifically to proteins like HspR or ParP when they are misfolded, a distinction we cannot yet quite make.

A PafE ring is formed of tightly packed α-helices and must be very rigid, which could impose a size limit on substrates. This observation is consistent with our earlier findings that disruption of *pafE* in *Mtb* does not result in accumulation of many proteins under normal culture conditions (11), and that the PafE proteome does not conspicuously overlap with the pupylome (30,31). While we do not yet know the function of ParP, it is likely to have ATPase activity and is not predicted to be essential for bacterial viability *in vitro* (20). Due to its robust interaction with PafE, a possibility is that ParP competes with other potential substrates and could perhaps be a regulator of PafE-mediated degradation *in vivo*. This hypothesis may be supported by our observation that the coproduction of HspR with PafE did not allow for their co-purification (data not shown), in contrast to what we observed for ParP and PafE.

Our studies have raised several fundamental questions that remain to be answered, including how does a substrate, either partially or totally disordered, get completely unfolded and threaded into a 20S CP for degradation in the absence of ATP? In addition, are other co-factors needed to help PafE target proteins to the proteasome? Finally, can PafE act as a protein chaperone for other substrates? Ongoing studies are working to answer these important questions.

## EXPERIMENTAL PROCEDURES

### Protein purification

All plasmids were made using PCR and sequenced by Genewiz, Inc (**Supplemental Table S1**). Coexpression of *pafE* and *parP* was achieved by insertion of a second ribosome-binding site and *parP* gene fragment after an XhoI site. A His_6_ tag was linked to C-terminus of ParP to co-purify the complex. Plasmids were transformed into *E. coli* BL21(DE3) and expression was induced by 0.2 mM IPTG at 16 °C. Bacteria were collected after over-night induction and homogenized in buffer containing 20 mM Tris·HCl (pH 8.0) and 300 mM NaCl. The cell lysates were loaded into a 5 mL Ni^2+^-nitrilotriacetate acid (Ni-NTA) agarose column and target proteins were eluted with a linear gradient concentration of imidazole. Further purification was performed using a Superose 6 10/300 GL gel-filtration column. To produce the fusion protein PafE_1-155_-PafE_1-174_, the gene fragment encoding PafE_1-155_ plus a C-terminal “GSGNS” linker (containing an EcoRI restriction site) was cloned into pET24b(+) (NdeI and NotI sites), followed by an insertion of the *pafE* gene between the EcoRI and NotI sites. The expression and purification methods for the fusion protein were the same as those for PafE alone. PafE variants, HspR, and 20S CPs were purified as described previously (5,13,32).

### In vitro degradation assay

For ParP degradation assays, 15 μg of purified WT 20S CPs were incubated at room temperature in reaction buffer containing 50 mM Tris·HCl (pH 8.0) and 5 mM MgCl_2_. The reaction was initiated by the addition of a four-fold molar excess of purified PafE:ParP or PafE_Δ155-174_:ParP. Reaction products were collected every 30 min and mixed with SDS protein sample buffer to stop the reaction. The degradation of ParP was assessed by SDS-PAGE. For the HspR degradation assay, 15 μg of purified WT 20S CPs was incubated with 10 μg WT or mutant PafE at room temperature in reaction buffer. The reaction was initiated by addition of a 30 μg HspR. Reaction products were collected every 30 min and mixed with SDS loading buffer to stop reaction. All degradation assays were done at least three times with essentially same results.

### Grid preparation and data collection

For cryo-EM analysis of PafE:20S CP complexes, 3 μl of purified sample of WT 20S CP complexed with PafE_Δ155-166_ were applied to a C-flat 2/1.2 holey carbon grid, which had been glow-discharged for 1 min by PELCO easiGlow (Ted Pella). Grids were plunge-frozen in liquid ethane using FEI Vitrobot with a blotting time of 7 s. Data were collected using an FEI Titan Krios 300 kV transmission electron microscope equipped with a Gatan K2 Summit direct detector. Automated data collection was carried out in super-resolution mode at a dose rate of 11.5 electrons/physical pixel/s. For each movie, 30 frames were acquired in 6 s with a total accumulated exposure of 58 electrons/Å^2^. A dataset of ~5,000 micrographs was collected using a defocus range between 1.0 and 3.0 μm.

For negative stain EM analysis of the PafE-PafE (PafE_1-155_-PafE_1-174_), ParP:PafE complexes, or PafE alone, 5 μl of protein sample were applied to a glow-discharged 300-mesh copper EM grid. The grid was then washed with water and stained twice for 45 s by adding 5 μl 2% (w/v) uranyl acetate solution. For the analysis of PafE-PafE, negatively stained EM grids were imaged in the Tecnai Spirit G2 BioTWIN transmission electron microscope operating at 120 kV. Micrographs were recorded in a dose of 12 electrons/Å^2^ at 30,000× magnifications in a Gatan Orius 830 CCD camera. For the analysis of ParP:PafE complex, the grid was loaded on a JEOL JEM-2010F transmission electron microscope operating at 200 kV equipped with a Gatan UltraScan 4000 CCD camera. Micrographs were recorded in low-dose mode (15 electrons/Å^2^) at 50,000× magnification (4,096 × 4,096 pixel), which corresponded to 2.12-Å/pixel sampling at the specimen level.

### Data processing and model building

For the negative-staining EM images of PafE-PafE (PafE_1-155_-PafE_1-174_), ParP:PafE complex, or PafE alone, particle selection and image processing were performed using EMAN2 software packages (33). PafE particles were selected in a semiautomatic manner with *e2boxer.py*. The particles were then subjected to 2D classification.

For cryo-EM analysis of PafE:20S CP, beam-induced motion correction and CTF estimation was done by MotionCor2 (34) and CTFFIND4 (35), respectively. Good micrographs were selected based on the extent and regularity of Thon rings. A total of 180,159 particles were automatically picked and applied to reference-free 2D classification using RELION2 (18). Classes with a clear density of 20S CPs, containing 141,275 particles, were selected for 3D classification. To make an initial reference, the EM structure of *Mtb* 20S CP with both ends capped with PafE_Δ155-166_ in our previous work was low-pass filtered to 50 Å. The particles were subsequently classified into eight classes without imposing any symmetry. Two out of the eight classes showed 20S CPs with one end capped with PafE. The two classes, containing 60,034 particles, were applied to high-resolution refinement without imposing any symmetry, which resulted in a map with an average resolution of 4.2 Å according to the FSC=0.143 criterion. Three classes containing 51,091 particles showed maps with good density of PafE on both ends of 20S CPs. Subsequent refinement with the D7 symmetry was performed using those particles. A mask was then generated to mask the 20S CP for high-resolution refinement using the D7 symmetry. Finally, a map of 20S CP upon PafE binding was obtained with an average resolution of 3.4 Å according to the FSC=0.143 criterion. Local resolution was estimated for the reconstruction with Resmap (36). Maps were visualized using UCSF Chimera (37).

The *Mtb* 20S crystal structure (PDB: 3MI0) was rigid body fitted into the 3.4-Å cryo-EM map and then iteratively refined using real-space refinement in Phenix (38). The resulted model was visually inspected in Coot (39). Several secondary structures and side chain rotamers were manually adjusted to best fit the density. The last four residues of PafE were also manually added to the extra density between the α subunits of the 20S CP. The quality of the final map was analyzed with Molprobity (40).

## Acknowledgments

Cryo-EM data was collected at the David Van Andel Advanced Cryo-Electron Microscopy Suite in the Van Andel Research Institute. K.H.D. thanks Chris Lima for helpful discussions. We thank Ashley Jordan for assistance with cloning. This work was supported by NIH grants AI070285 (to H.L.), AI088075 (to K.H.D), and T32 AI007180 and F30 AI110067 (to J.B.J), and the Van Andel Research Institute (to H.L.). K.H.D. holds an Investigator in the Pathogenesis of Infectious Disease Award from the Burroughs Wellcome Fund. We have no conflict of interest to declare.

## Accession codes

The doubly PafE-capped and the singly PafE-capped *Mtb* 20 CP cryo-EM 3D maps have been deposited in the EMDB database with accession codes EMD-7097 and EMD-7098, respectively. Their corresponding atomic models have been deposited in the RCSB PDB with accession codes 6GBL and 6GBO, respectively.

## Author contributions

K.H. K.H.D, and H.L. designed the research. K.H. J.B.J, S.Z., and A.K. performed biochemical experiments. K.H. purified proteins and did negative stain EM, and K.H and G.Z did cryo-EM. K.H., K.H.D, and H.L. analyzed the data and wrote the paper. All authors contributed to finalizing the manuscript and approved the final version of the manuscript.

## Additional information

Supplementary information is available in the online version of the paper.

## Supplemental Information

**Table S1.** List of plasmids and strains used to produce proteins in this study.

| Proteins | Plasmids/Host | Relevant genotype (Reference) |
| --- | --- | --- |
| PafE | pET24b(+)/BL21(DE3) | Kan<sup>r</sup>, N-terminal His<sub>6</sub>-tag (1) |
| PafE<sub>Δ155-166</sub> | pET24b(+)/BL21(DE3) | Kan<sup>r</sup>, N-terminal His<sub>6</sub>-tag (2) |
| PafE<sub>Δ155-174</sub> | pET24b(+)/BL21(DE3) | Kan<sup>r</sup>, N-terminal His<sub>6</sub>-tag |
| PafE<sub>P65G</sub> | pET24b(+)/BL21(DE3) | Kan<sup>r</sup>, N-terminal His<sub>6</sub>-tag |
| PafE<sub>F138E</sub> | pET24b(+)/BL21(DE3) | Kan<sup>r</sup>, N-terminal His<sub>6</sub>-tag |
| PafE<sub>1-155</sub>-GSGNS-PafE<sub>1-174</sub> | pET24b(+)/BL21(DE3) | Kan<sup>r</sup>, PafE<sub>1-155</sub> N-terminal His<sub>6</sub>-tag, WT PafE |
| PafE:ParP | pET32a(+)/BL21(DE3) | Amp<sup>r</sup>, WT *pafE*, *parP*-his<sub>6</sub> |
| PafE<sub>1-155</sub>:ParP | pET32a(+)/BL21(DE3) | Amp<sup>r</sup>, *pafE*<sub>1-155</sub>, *parP*-his<sub>6</sub> |
| HspR | pET32a(+)/BL21(DE3) | Amp<sup>r</sup>, *hspR*-his<sub>6</sub> (1) |
| PrcB, PrcA | pET32a(+)/BL21(DE3) | Amp<sup>r</sup>, *prcA*, *prcB*-his<sub>6</sub> (1) |
| PrcB, PrcA<sub>R26A</sub> | pET32a(+)/BL21(DE3) | Amp<sup>r</sup>, *prcA*<sub>R26A</sub>, *prcB*-his<sub>6</sub> |
| PrcB, PrcA<sub>K52A</sub> | pET32a(+)/BL21(DE3) | Amp<sup>r</sup>, *prcA*<sub>K52A</sub>, *prcB*-his<sub>6</sub> (2) |

**Figure S1.**
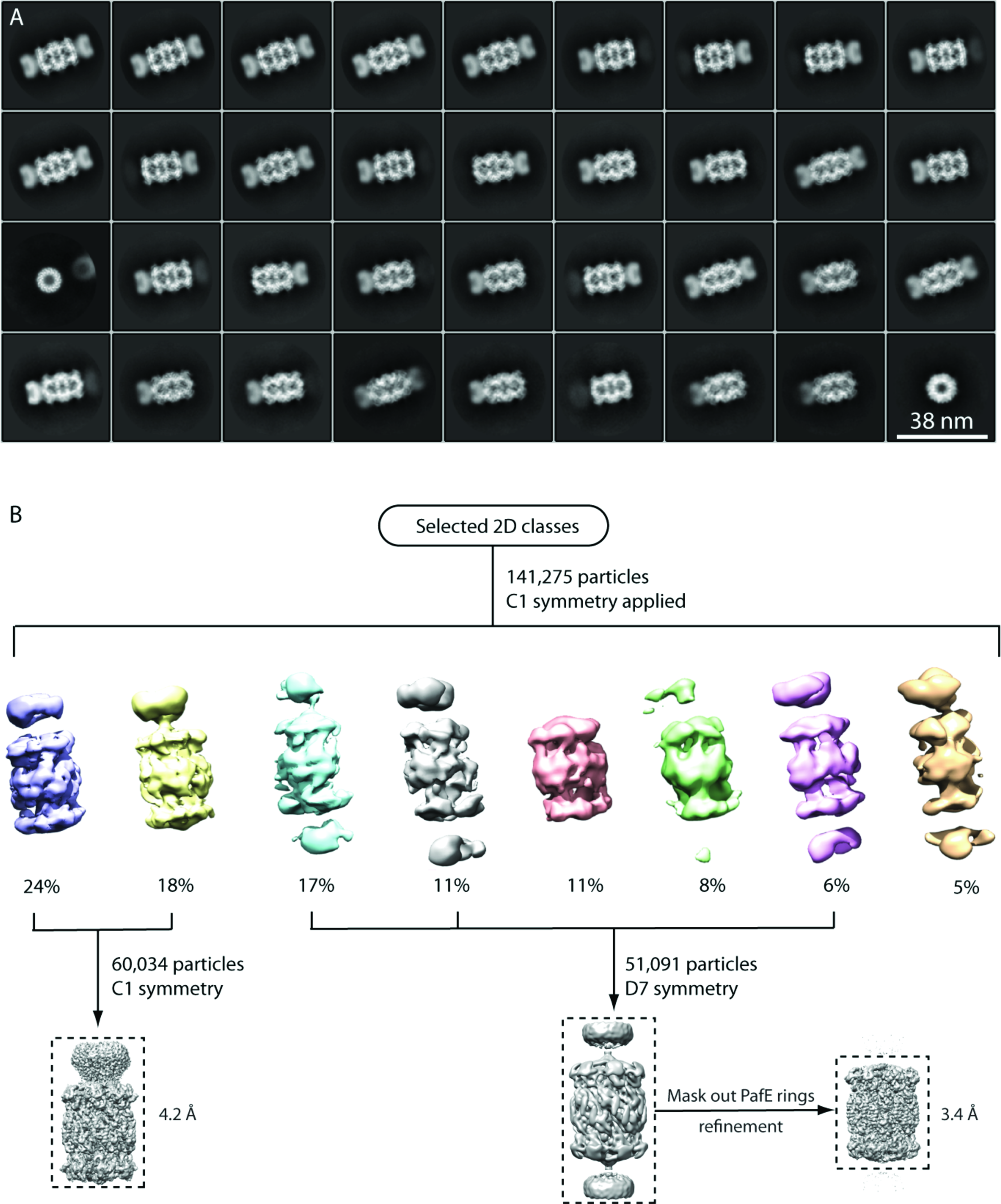
Cryo-EM studies of PafE_Δ155-166_:20S CP complex. (**A**) Representative 2D class averages of roughly 180,000 auto-picked particles. (**B**) The procedures outline to generate the 3D reconstructions of PafE_Δ155-166_:20S CP complex. The eight classes obtained from 3D classification are showed and the distribution is indicated by percentage below. The classes for high-resolution reconstructions are indicated. The dashed lines show the mask used in refinement.

**Figure S2.**
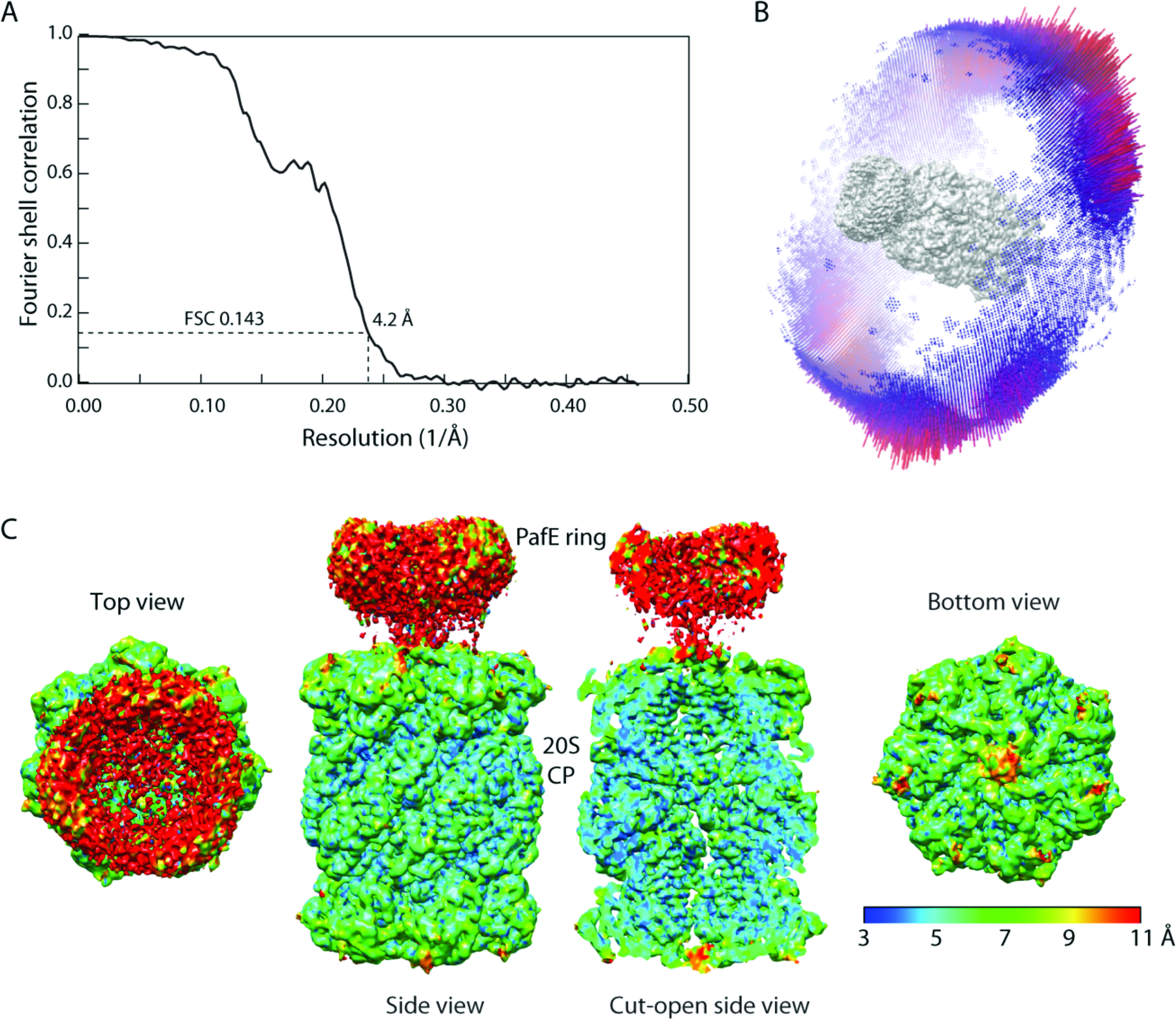
Fourier shell correlation and local resolution estimate for EM reconstruction of one-end-capped 20S CP. (**A**) Fourier shell correlation curve of 20S CP with PafE_Δ155-166_ bound at one end. No symmetry was imposed during the 3D reconstruction. (**B**) Euler angle distribution of the particles used in the reconstruction. (**C**) Surface representation of the EM map colored according to the local resolution estimate by ResMap.

**Figure S3.**
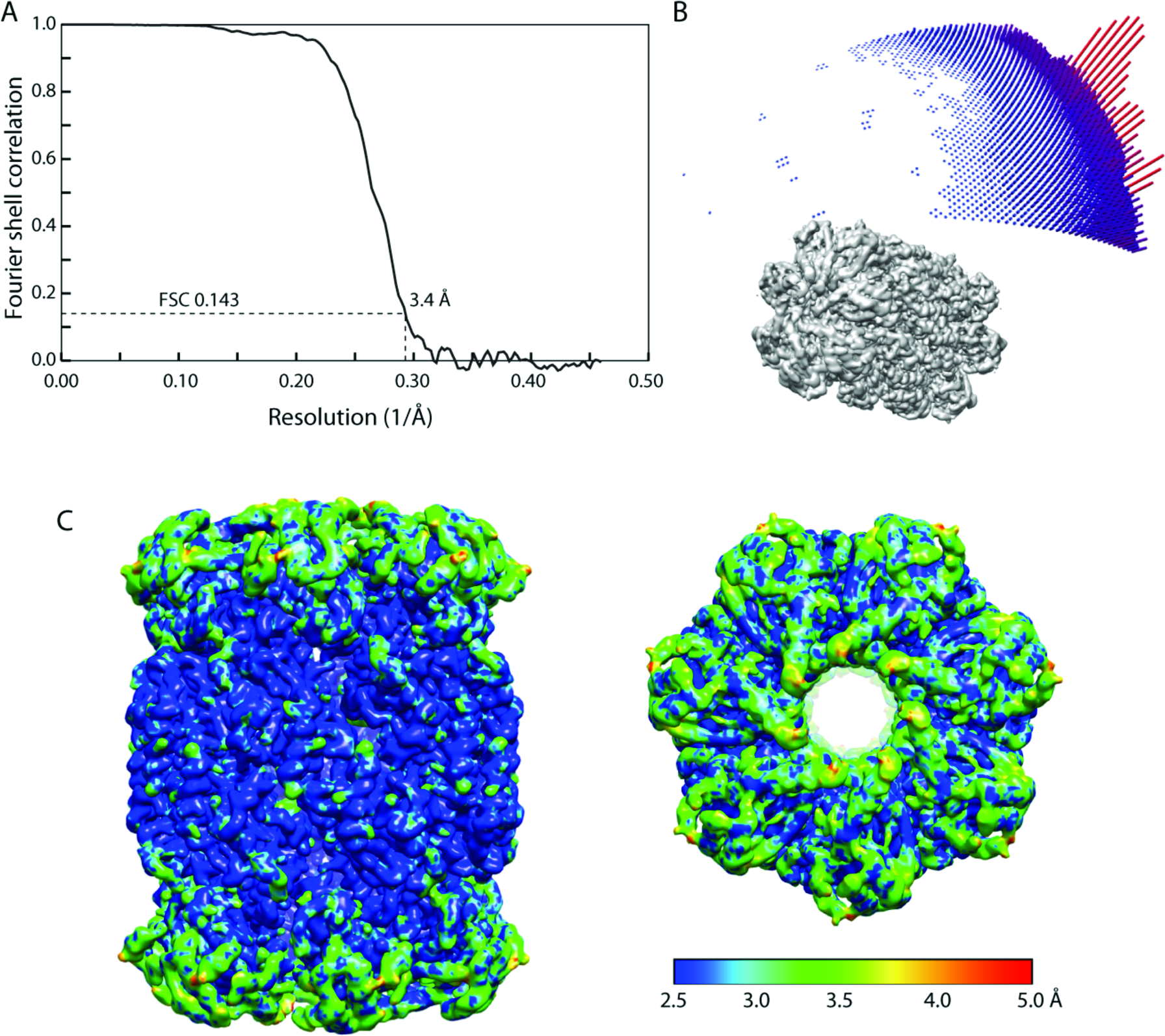
Fourier shell correlation and local resolution estimate for EM reconstruction of two-ends-capped 20S CP. (**A**) Fourier shell correlation curve of 20S CPs with PafE_Δ155-166_ bound at both ends. The PafE regions were masked out during final refinement, and the D7 symmetry was imposed in 3D reconstruction. (**B**) Euler angle distribution of the particles used in the reconstruction. (**C**) Surface representation of the EM map colored according to the local resolution estimate by ResMap.

**Figure S4.**
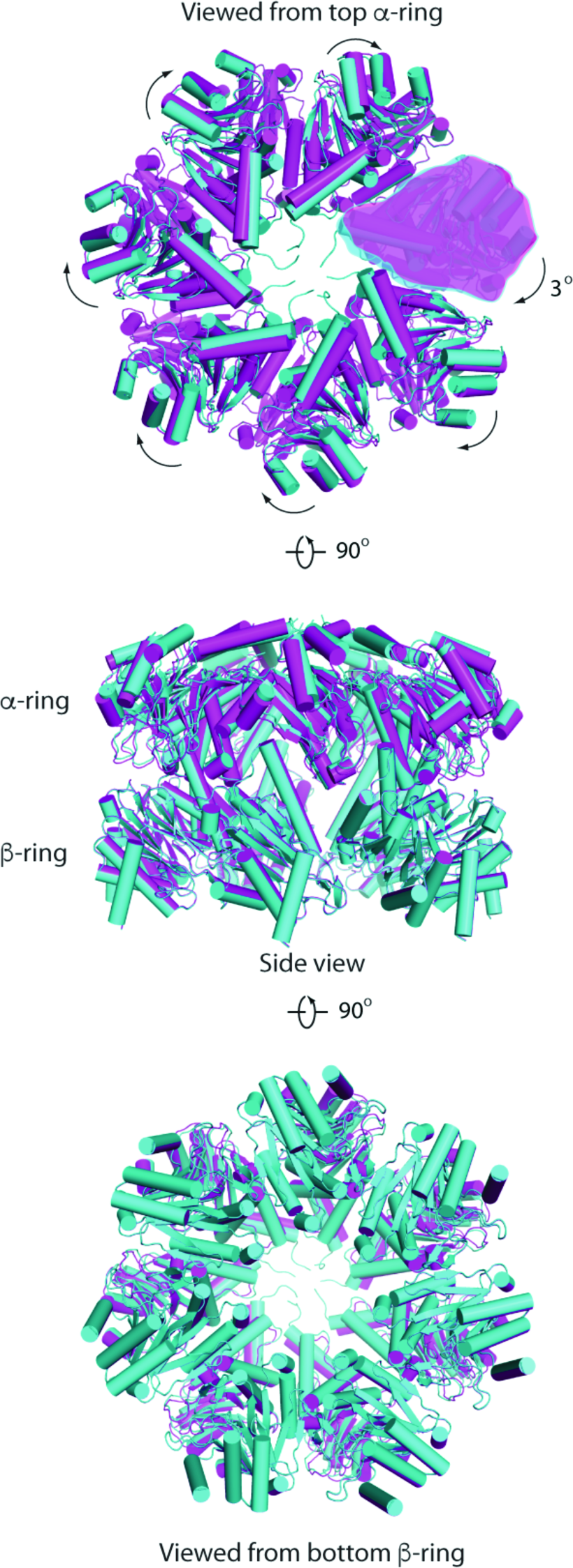
Alignment of PafE-capped vs. uncapped half 20S CPs. The PafE-capped 20S CP model in Figure 2 was divided into two halves. The PafE-capped and uncapped halves were colored in magenta and cyan, respectively. The top, side, and bottom views are shown in the upper, middle, and lower panels.

**Figure S5.**
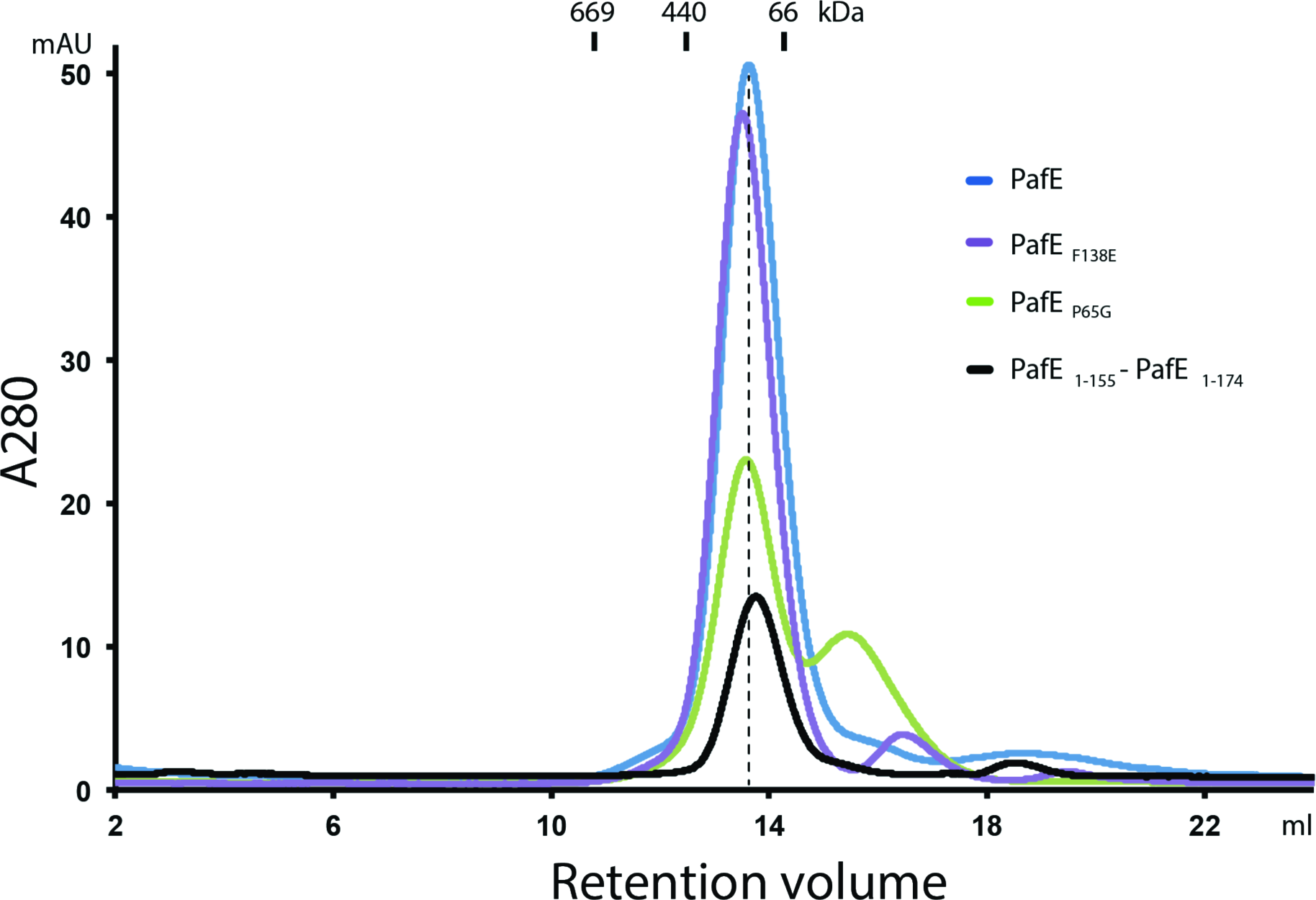
Size exclusion chromatography profiles of wild type PafE and mutant alleles. Superose 6 10/300 GL gel filtration column is used for the chromatographic studies. The curves of PafE, PafE_F138E_, PafE_P65G_, and PafE_1.155_-PafE_1.174_ are colored in blue, purple, green, and black, respectively. Size standards are marked beneath the chromatographic panel.

